# Structural brain connectivity predicts acute pain after mild traumatic brain injury

**DOI:** 10.1101/2021.11.12.468345

**Authors:** Paulo Branco, Noam Bosak, Jannis Bielefeld, Olivia Cong, Yelena Granovsky, Itamar Kahn, David Yarnitsky, A. Vania Apkarian

## Abstract

Mild traumatic brain injury, mTBI, is a leading cause of disability worldwide, with acute pain manifesting as one of its most debilitating symptoms. Understanding acute post-injury pain is important since it is a strong predictor of long-term outcomes. In this study, we imaged the brains of 172 patients with mTBI, following a motorized vehicle collision and used a machine learning approach to extract white matter structural and resting state fMRI functional connectivity measures to predict acute pain. Stronger white matter tracts within the sensorimotor, thalamic-cortical, and default-mode systems predicted 20% of the variance in pain severity within 72 hours of the injury. This result generalized in two independent groups: 39 mTBI patients and 13 mTBI patients without whiplash symptoms. White matter measures collected at 6-months after the collision still predicted mTBI pain at that timepoint (n = 36). These white-matter connections were associated with two nociceptive psychophysical outcomes tested at a remote body site – namely conditioned pain modulation and magnitude of suprathreshold pain–, and with pain sensitivity questionnaire scores. Our validated findings demonstrate a stable white-matter network, the properties of which determine a significant amount of pain experienced after acute injury, pinpointing a circuitry engaged in the transformation and amplification of nociceptive inputs to pain perception.

## Introduction

Traumatic brain injury is a leading cause of death and disability in the United States(1), with important socioeconomic costs. Motor vehicle collisions are a frequent cause of mild TBI (mTBI), which often is accompanied with whiplash associated disorders (WAD) given the quick shift of forces during a crash and the consequent abrupt and brisk back-and-forth movement of the neck. These patients usually present with a range of clinical symptoms including stiffness, dizziness, nausea, and mental confusion, but the most frequent and debilitating clinical manifestation is pain in the head/neck area(2), which may last from days to weeks and – for a large proportion of patients – can become chronic(3). Managing pain post-injury is of utmost importance since it reflects on the patients’ recovery process: mTBI patients with acute pain-related disabilities are at higher risk for chronic pain(3) and are more likely to develop clinically significant anxiety, depression, sleep disturbances, and post-traumatic stress disorder(4)

Early acute pain after mTBI is, however, poorly understood. The etiology of pain does not map well onto injury-related imaging findings(5),(6),(7) and is only partially explained by psychological and psychophysical pain characteristics(8,9). It is known that psychological factors play a role in early acute pain, with state anxiety, depression, and pain catastrophizing modestly explaining additional variance in pain severity(10). At the chronic stage, it is also known that these patients have altered psychophysical pain indices, including hyperalgesia to painful stimuli, inefficient conditioned pain modulation, and enhanced temporal summation of pain(11). While it is thought that both pro-nociceptive and affective/emotional processes are driven by brain-centric mechanisms, so far, no studies have examined the brains of these patients in relation to acute mTBI pain. Consequently, how the central nervous system reacts to, is changed by, or can predispose someone to experience pain after acute mTBI injury is unknown. Peering into the subject’s brains and exploring the mechanisms underlying pain at the early acute stage is a promising approach to resolve the relative contribution of psychological, psychophysical, and nociceptive processes in post-injury pain and can shed light on the mechanisms that underlie pain perception immediately after an emotionally charged, pain-inducing incident.

In this study, we used MRI to study brain functional and structural properties of mTBI patients after a motorized vehicle accident and probed for properties that can determine or predispose patients to experience pain after injury. Using a machine-learning approach, we examined the functional and structural brain networks associated with early, acute pain. This study sheds light on the properties and emergence of pain, particularly after injury, with implications for the treatment and management of mTBI.

## Results

### Predicting pain after mTBI using functional and structural connectivity

We used a machine-learning approach to predict pain from resting-state functional magnetic resonance imaging (rsfMRI) and diffusion tensor imaging (DTI) brain connectivity. Discovery (70% of the data, N_rsfMRI_ = 94 and N_DTI_ = 88 after outlier exclusion) and hold-out (N_rsfMRI_ = 43 and N_DTI_ = 37 after outlier exclusion) datasets were used to build the model and assess generalizability (see Fig. 1 and methods for an explanation of the machine-learning pipeline). For structural connectivity, after 10-folds cross-validation (CV), the best *p* threshold for univariate feature selection was determined to be *p* = 0.01 (negative features model, highest CV *r*_*predvsactual*_ = .33; positive model *n*.*s*.), leading to the selection of 26.5 features on average, which mostly appeared inconsistently across CV folds (see Supplementary Fig. 1).

**Figure 1.**
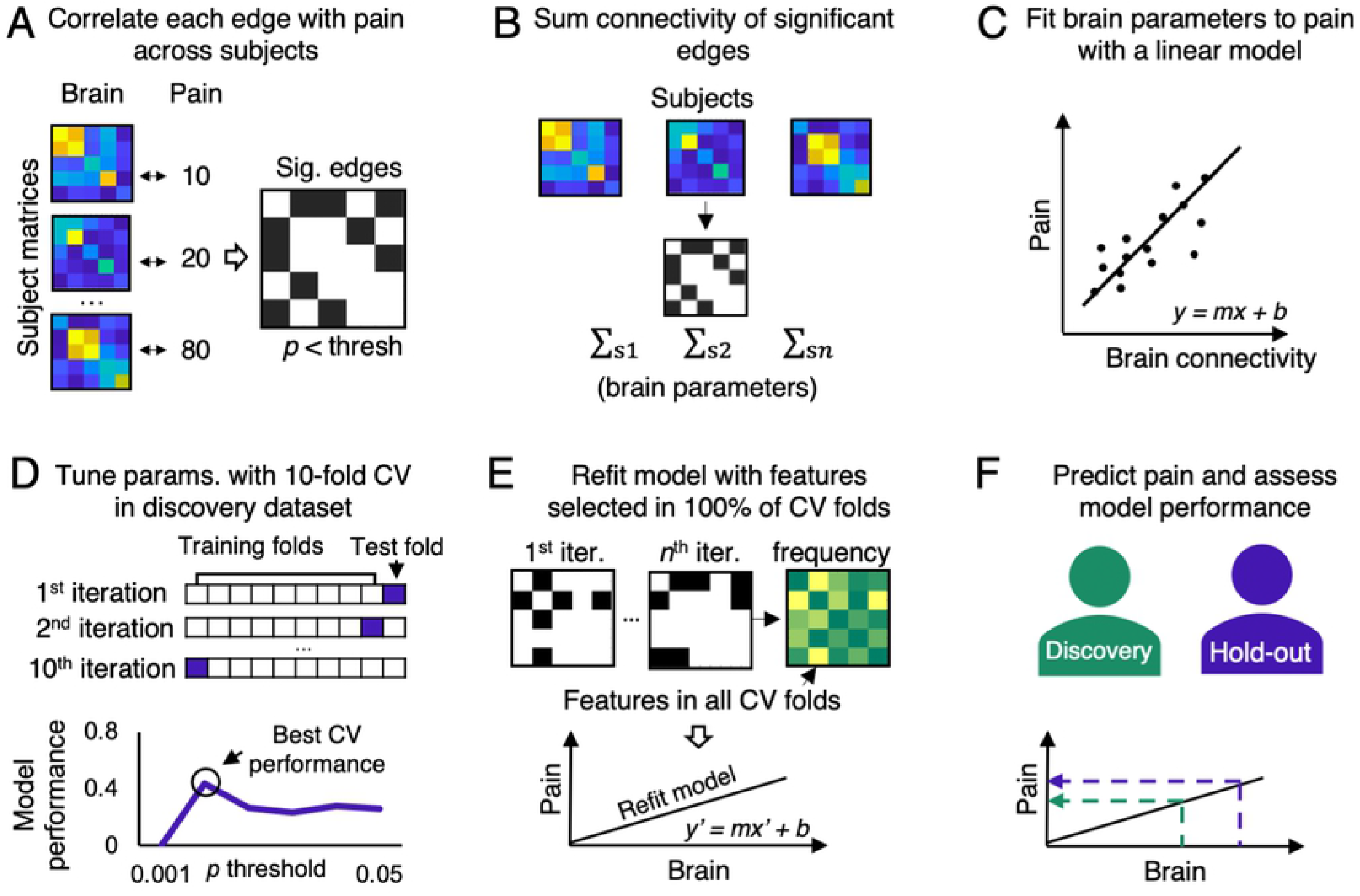
Machine-learning pipeline to identify mTBI acute pain related brain circuitry. We used Connectome-based Predictive Modelling approach to model large-scale connectivity. Upper panel, modelling approach: **A**. First, all pairs of connectivity within the 105 ROIs were correlated with pain (across-subjects) to select a series of significant edges (feature selection step). **B**. All significant edges were then summed into a single value (separately, for positive and negatively correlated edges, data-reduction step). **C**. This summarized pain model was then regressed against pain to obtain a linear equation predicting pain, which was tested in out-of-sample subjects. Lower-panel, validation approach: the above approach was embedded into a machine-learning algorithm that attempts to generalize the model impartially. **D**. The total sample here was separated into a discovery (70%) and a hold-out datasets (30%). Within the discovery dataset, cross-validation was conducted to impartially determine the *p*-threshold used in the feature selection step. **E**. To improve model stability, after the best *p*-value was determined, a new model was built based only on the features that were significant in all the CV folds (features occurring in 100% of the 10-fold CV iterations). **F**. Model performance was assessed within-sample by applying the linear equation to the discovery dataset and out-of-sample by applying the linear equation in the hold-out dataset to assess generalization.

Only three white-matter (WM) tracts were selected on all cross-validation folds: tract 1, left Precentral gyrus - left Postcentral Gyrus; tract 2, left Thalamus - left Superior Parietal Lobule; and tract 3, right Planum Polare - left Superior Lateral Occipital Cortex (Fig. 2A). Within the discovery dataset, these three features, together, show an *r*_*predvsactual*_ = 0.47 [95% CI: 0.324, 0.572], and *p* < .001 (Fig. 2B). By applying the linear model built on the discovery dataset, we were able to predict patient’s pain in the hold-out dataset (Fig. 2B): the model showed a *r*_*predvsactual*_ = 0.43 [95% CI: 0.158, 0.593], *p* = .007, thus suggesting generalizability. We further examined if the WM connectivity model could predict pain in the sub-sample of mTBI patients without whiplash symptoms (WAD score = 0, N = 13). Since these subjects were not used in previous analyses, they represent a second independent group on which generalizability can be assessed. The model was again successful at predicting patients’ pain and had a *r*_*predvsactual*_ = .53 [95% CI = 0.058, 0.697], see Fig. 2B, although this correlation did not reach significance (*p* =.061), potentially due to the small sample size. Meta-analytical prediction metrics were obtained by assessing the pooled prediction performance in the three datasets (N = 138). Results replicate the previous findings, with a *r*_*predvsactual*_ = .44 [95% CI = 0.320, 0.532], *p* < .001, see Fig. 2C. For the three tracts, WM connectivity was significantly correlated with pain ratings (all *r*s > -0.3, *p*s < .001), with higher connectivity strengths leading to lower observed pain (Fig 2D). Finally, to discard the possibility that the WM connectivity strength is associated with the severity of the clinical manifestation we explicitly compared the connectivity strength of the three tracts between patients with a WAD score of 0, 1 and 2, reflecting whiplash signs and symptoms severity. A one-way ANOVA showed no significant differences in connectivity between groups (*F*(2,135) = 1.11, *p* = .33, Fig. 2E).

**Figure 2.**
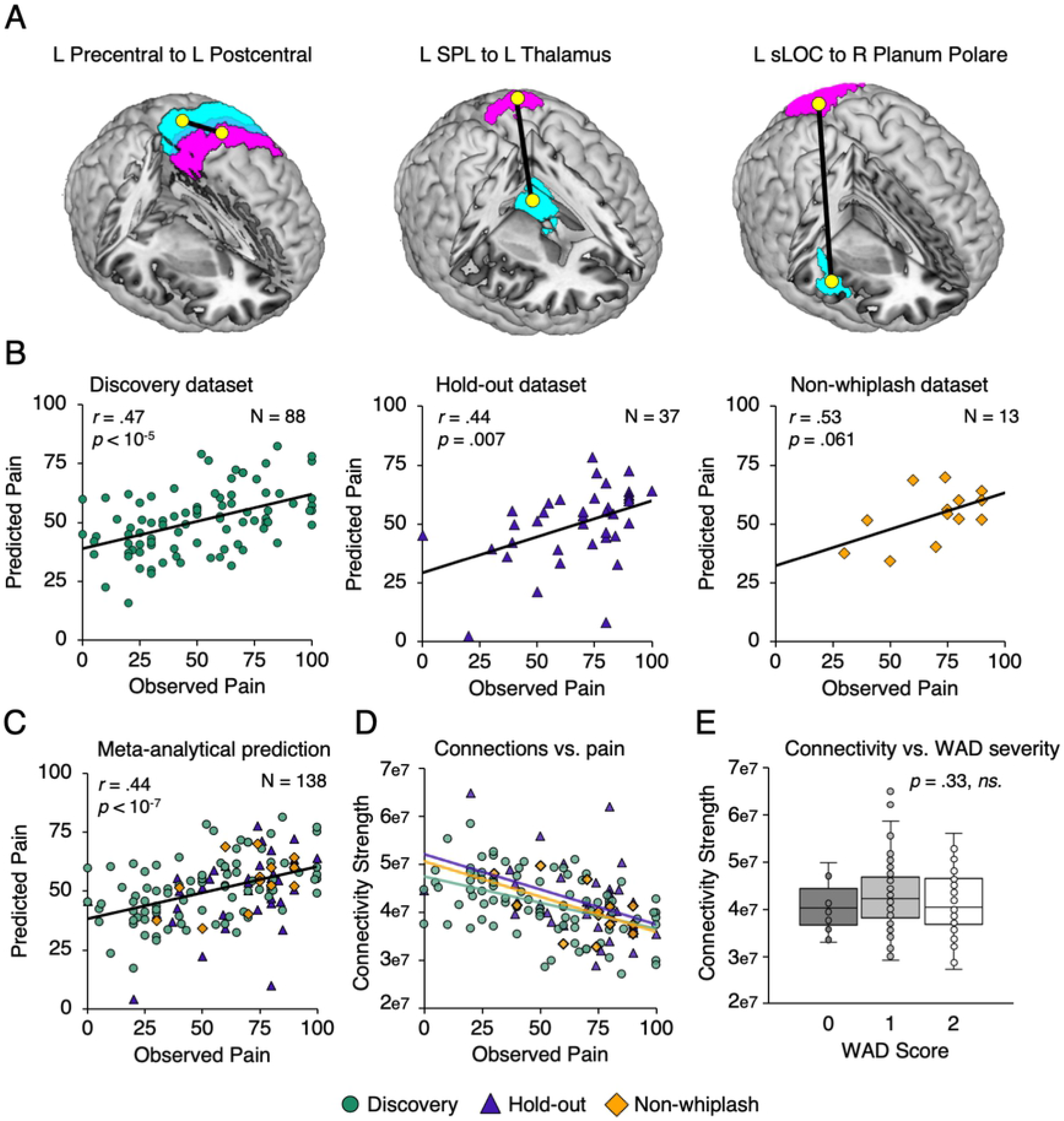
Structural connections between three pairs of cortical regions reflect acute mTBI pain. Three features were selected by the machine-learning algorithm **(A)**, and a model was cast. The model’s predicted values were significantly correlated with actual pain in the discovery dataset **(B, left)** and in the hold-out dataset **(B, middle)** with equivalent effect sizes. The model was also able to predict pain in a third independent group of mTBI patients with no whiplash-like symptoms **(B, right)**. Meta-analytical predictive performance metrics were calculated to summarize the results from the three groups **(C)**, showing again reliable associations between predicted and observed pain. **(D)** Raw connectivity strength and their association with pain are illustrated, showing a negative association whereby more connections lead to less reported pain. **(E)** Connectivity strength of the three tracts compared between subjects with WAD scores of 0, 1, and 2 show no significant differences, supporting no association between symptom severity and connectivity strength. left superior parietal Lobule, L SPL; left superior lateral occipital cortex, sLOC.

For rsfMRI analyses, no statistically significant model emerged for either the positive or negative features model at any threshold (see Supplementary Fig. 2). Hence, we conclude that using this approach, rsfMRI is unable to predict early acute pain. Further analyses are reported for structural analyses only.

### Model predictions do not depend on age, sex, total intracranial volume

We next asked whether this finding could be explained by confounds, such as age, sex, and total intracranial volume (TIV). We conducted a sequential linear regression on the discovery dataset with age, sex, and TIV as independent variables and pain as the dependent variable. This model was not significant (*p* = .074) and only explained 4.6% of the variance in pain ratings. Adding the structural connectivity score to this model resulted in a significant model that explained 22% of the variance in the pain ratings (*p* < .001); this increase in explained variance was statistically significant (model 1 *vs*. model 2, *F*[1, 83] = 20.29, *p* < .001).

### Spatial specificity within thalamic and somatosensory networks

Given that the sensorimotor network follows a somatotopic organization and the thalamus has well-characterized nuclei implicated in multiple sensory and affective processes, we further tested the spatial specificity of both regions. Thus, brain regions from the left precentral and postcentral gyrus were parcellated based on their body-part representation (face, arm, trunk, leg) using masks published previously(12) (Fig. 3A). The connectivity between each pair of regions was estimated and correlated with baseline pain. The number of connections between the precentral-postcentral face (*r* = -.26, *p* = .004, Fig. 3A), the precentral-postcentral arm area (*r* = -.24 *p* = .008), and precentral arm and postcentral face (r = -.21, *p* = .019), were significantly associated with pain ratings.

**Figure 3.**
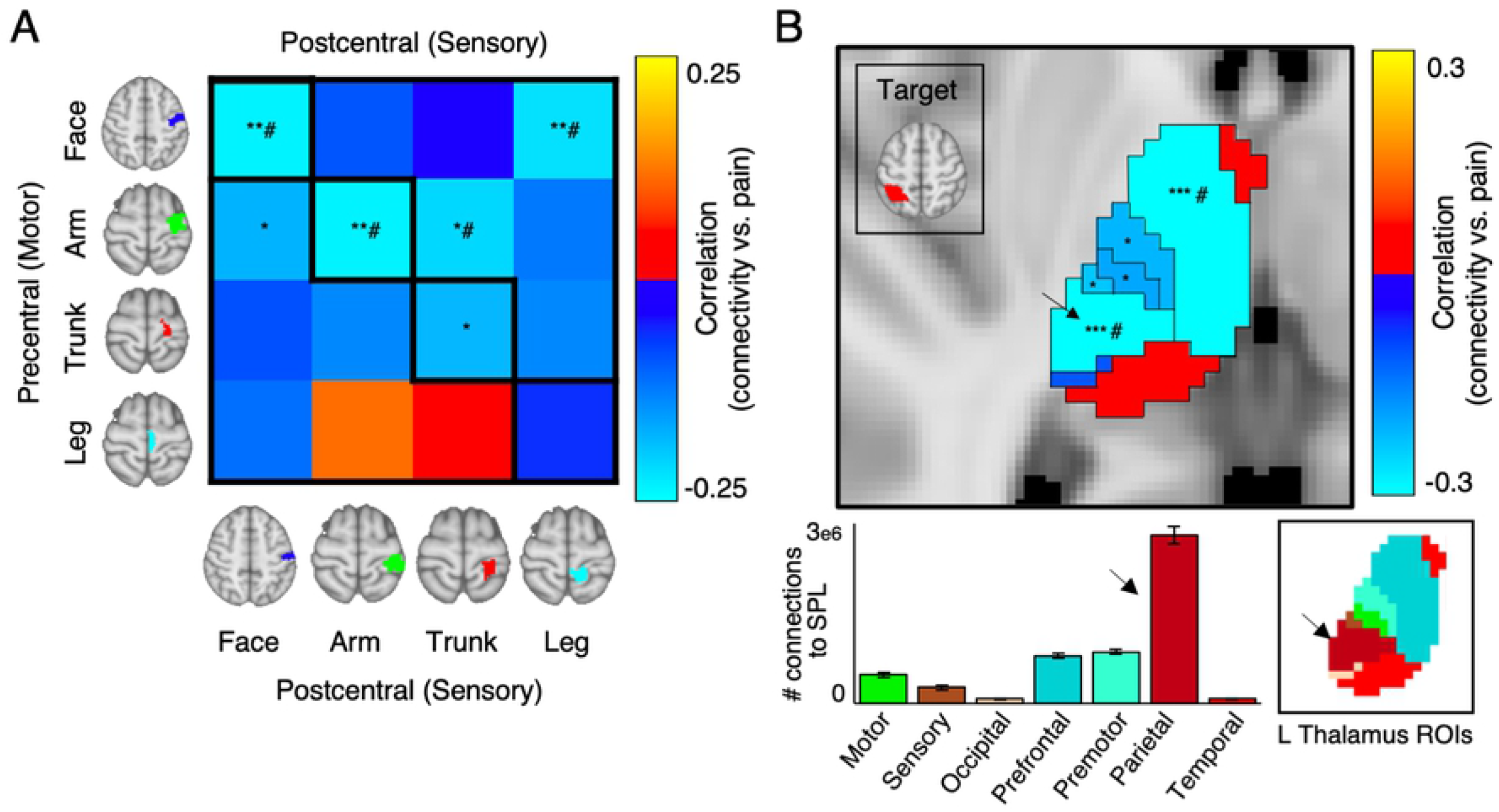
Somatotopic organization of structural connectivity within the sensorimotor cortex and the thalamocortical system reveal topographically appropriate linkages related to acute mTBI pain. Post-hoc analysis looking at left PreC-PostC and left Thal-SPL connectivity at a higher granularity. (**A)** Within the sensorimotor homunculi, the most predictive connectivity is between the face and arm ROIs. Connectivity between the precentral arm and postcentral trunk (site of injury, i.e. head/neck representation in the motor and sensory homunculi, respectively) also show significant correlation with observed pain. (**B)** For the thalamus, the strongest predictor was the VPL nuclei (top panel), together with an anterior/medial region. The VPL nuclei was also the nucleus with strongest absolute connectivity to the parietal cortex (bottom panel). * *p* < .05; ** *p* < .01; # survives FDR correction for multiple comparisons. Thalamus ROIs can be depicted in the bottom right panel.

We also examined the left thalamus using the FSL thalamic connectivity atlas(13), containing seven regions of interest (ROIs), each highlighting connectivity to different brain areas (prefrontal, premotor, motor, sensory, parietal, occipital, temporal). The connectivity between each thalamic ROI and the left Superior Parietal Lobule (SPL) was calculated and correlated with reported pain. We found evidence of spatial specificity – the thalamic nucleus most predictive of pain was the subregion mostly connected to the Parietal cortex, corresponding spatially to the posterior lateral thalamus area (*r* = -.30, *p* < .001, Fig. 3B); further, we also observed strong associations between the connectivity of the prefrontal ROI and left SPL, tapping into more anterior-medial thalamus (*r* = -.30, *p* < .001). We further observed significant correlations between lateral thalamic nuclei connecting to sensory (*p* = .009), motor, and premotor areas (*p* = 0.027, and *p* =.017, respectively). The other thalamic ROIs (connected to the temporal and occipital cortex) were not significantly associated with pain.

### Predictive brain parameters are associated with psychophysical indices of pain sensitivity

To ground these findings in clinical properties, we tested for an association between brain, psychophysical and clinical parameters. We found significant positive associations between WM connectivity strength of all tracts and the conditioned pain modulation (CPM, *r* = .21, *p* = .019), the temperature at which subjects rated a pain of 50/100 (suprathreshold pain, termed Pain50(8), *r* = .20, *p* =.027), as well as negative associations with scores in the pain sensitivity questionnaire (PSQ, *r* = -.19, *p =* .038). Individually, pressure-pain CPM response was mostly associated with precentral-postcentral connectivity (*r* = .25, *p* = .006), while connectivity between the right Planum Polare and the superior Lateral Occipital Cortex was associated with the perceived stress scale (*r* = .20, *p* = .034). Other clinical and psychological parameters did not show significant results (Fig. 4A).

**Figure 4.**
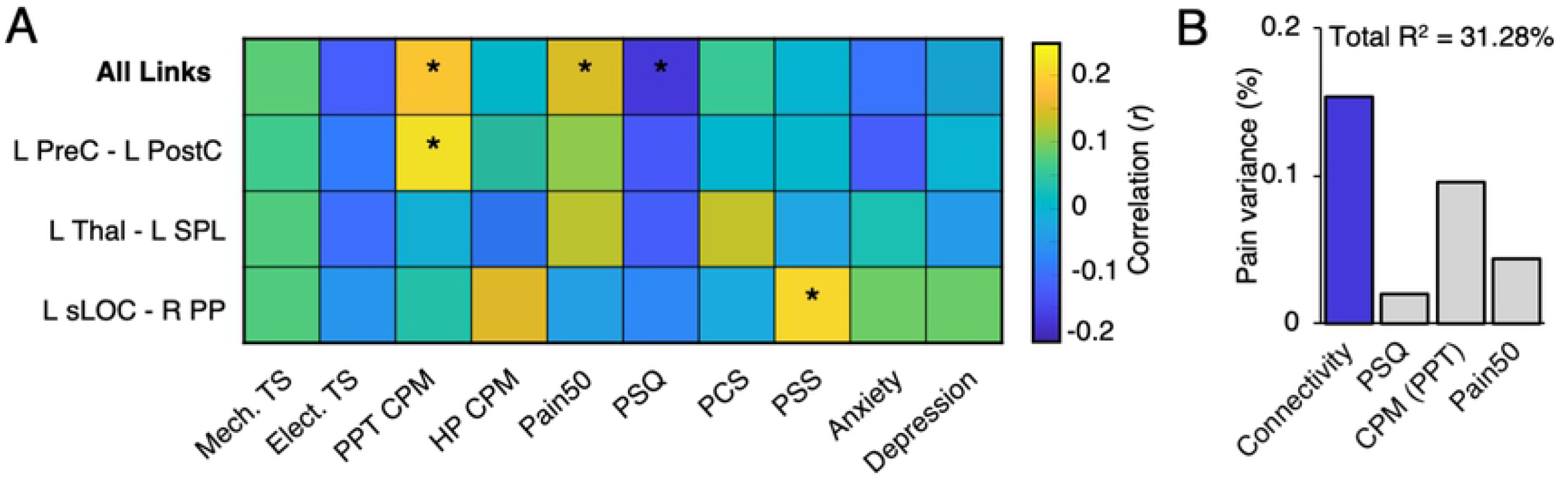
A multi-parameter model for predicting acute and longer-term mTBI pain. **(A)** Correlations between the predictive structural connectivity tracts and psychophysical-psychological variables. Higher pressure-pain conditioned pain modulation (CPM) and suprathreshold pain (Pain50) scores were associated with stronger connectivity, and lower pain sensitivity questionnaire (PSQ) scores were associated with stronger connectivity. Individually, left PreC-PostC was mostly associated with CPM, while R PP – sLOC was associated with perceived stress scale (PSS). Mechanical (Mech. TS) and electrical (Elect. TS) temporal summation, Heat-pain conditioned pain modulation (HP CPM), Pain Catastrophizing (PCS), Anxiety and Depression did not show any significant associations. **(B)** Relative importance metrics for connectivity, PSQ, CPM, and Pain50, when predicting pain. A model with the four parameters (Brain, psychological, and psychophysical parameters) explains 31% of the variance in pain, with connectivity parameters accounting for 50% of the full model variance (i.e. 15% of unique variance in observed pain).

Given that these clinical parameters have been previously associated with pain ratings in these individuals(8,9), we further wanted to disentangle the relative contribution of these psychological and psychophysical dimensions to that of the brain. To do so, we performed a relative importance analysis (using the R package *relaimpo*(14)). While the total model with the four parameters (CPM, Pain50, PSQ, and Brain Connectivity) explained 33.6% of the variance in pain ratings (Fig. 4B), brain connectivity alone explained 15% unique variance in pain ratings.

### The predictive ability of the model is not contingent on machine-learning pipeline

In order to ensure these results were not contingent on the modelling approach, we repeated the analyses using a simple linear regression combining both positively and negatively associated tracts and without summing the tracts into a single value. After cross-validation, the best performing threshold for feature selection was *p* = .005 (CV *r*_*predvsactual*_ = .15), leading to two highly reliable features (tract 1, left Precentral gyrus - left Postcentral Gyrus; tract 2, left Thalamus - left Superior Parietal Lobule; tracts 1 and 2 from the previous analyses). Within the discovery dataset, the model obtained an *r*_*predvsactual*_ = 0.37 [95% CI: 0.192, 0.503], *p* < .001. Within the hold-out dataset, this model obtained a *r*_*predvsactual*_ = .40 [95% CI: 0.115, 0.577], *p* = 0.014. Unsurprisingly, predictions and statistical results from machine-learning and linear regression models were similar, with the correlation of predicted pain values from both methods being *r* = .90 and .96 for the discovery and hold-out datasets, respectively.

### Structural connectivity parameters are stable over time, and explain future pain

Since this study is part of a larger project aimed at discovering brain predictors of transition into chronic pain in mTBI patients, we examined the pain-ratings of these patients over time. The strength of WM connectivity at baseline was associated with patients’ pain one, three, and six months after baseline (all *ps* < .05, Fig. 4E). This effect, however, was substantially reduced when controlling for pain ratings at baseline. We then assessed whether these connectivity indices were static in time; hence, we examined DTI data from a sub-sample of these patients that repeated the MRI protocol at six months (N = 36) and one year (N = 13) after injury. We compared, within-subject, whether the connectivity strength of the three tracts changed over time. Paired t-tests show that the connectivity does not change over time (*p* = .74, *p* = .52, 6 months and 1 year, respectively, Fig.5A). Importantly, DTI parameters collected at six months post-injury were also significantly associated with pain at six months (*r* = -41, *p* = .012, see Fig. 5B). This result, however, did not hold at one year after injury (*r* = .25, *p* = .4), despite DTI parameters not changing and a significant proportion of patients (50%) still reporting pain.

**Figure 5.**
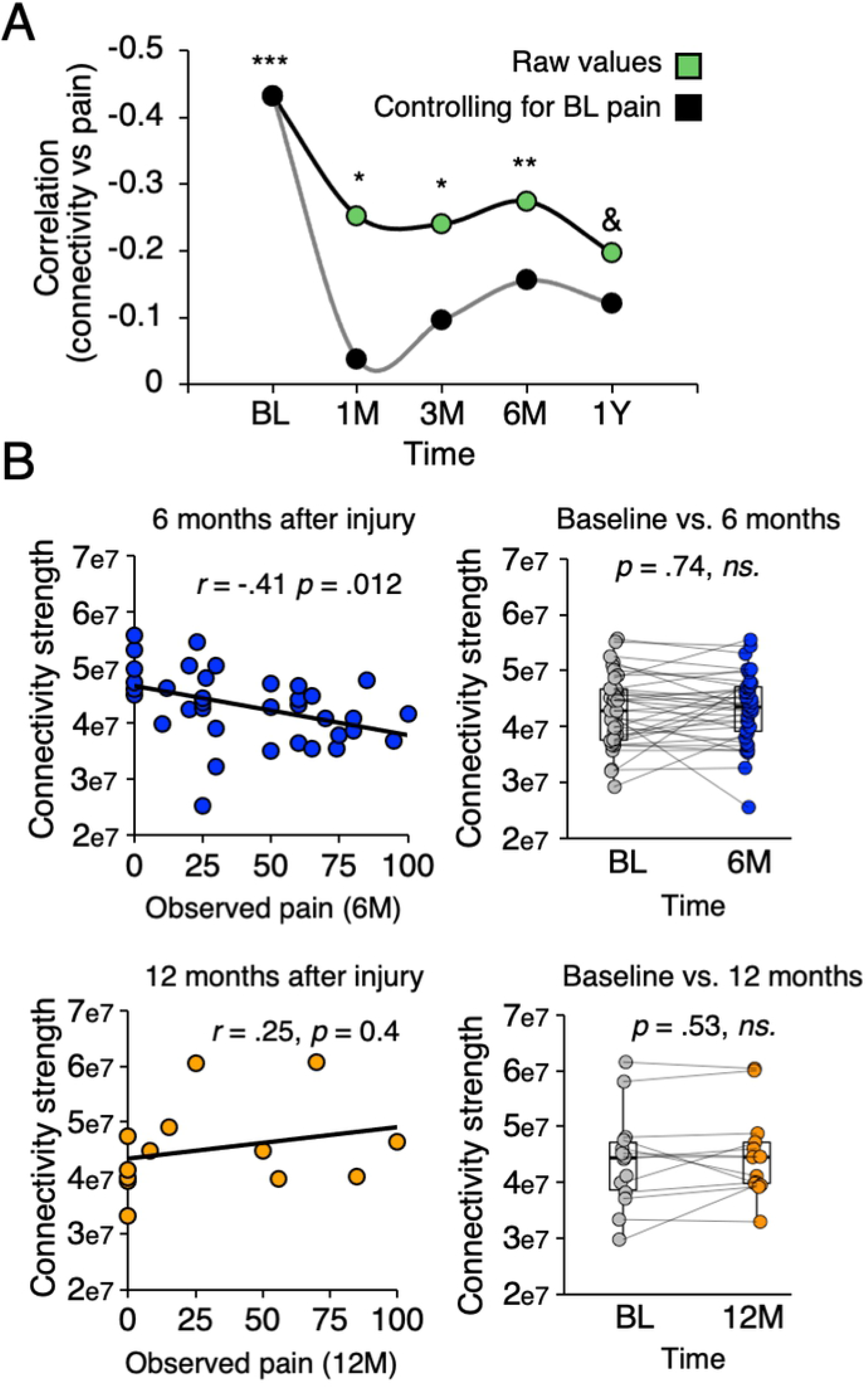
Stability of structural connectivity related to acute mTBI pain over 12 months and their relationship to long-term mTBI pain. **(A)**. Model performance at predicting pain over time; baseline parameters are associated with observed pain at baseline (BL), 1 month (1M), 3 months (3M), 6 months (6M) and a year (1Y, marginally significant) after a motorized vehicle crash, but not when controlling for baseline pain. **(B, left panel)** connectivity strength of the three predictive features measured at 6 months post-MVC is still significantly associated with reported pain at 6 months, but not at 12 months. **(B, right panel)** connectivity remains stable from baseline to 6 months, and after one year. Lines represent paired differences within subject. * *p* < .05; ** *p* < .01; *** *p* < .001; & *p* = .051.

## Discussion

In this study, we examined the brains of mTBI patients suffering from early acute pain, attempting to characterize brain networks underpinning acute pain after a motorized vehicle collision injury. Our results can be summarized in four key points. First, we show that white-matter brain properties – but not functional properties – explain a sizable variance of the pain after mTBI, hence predisposing patients to report more pain after acute injury. Second, we demonstrate that these findings are not dependent on injury-related, clinical, or demographic characteristics of the patients. Third, these white-matter connections map well onto physiological-psychological characteristics, namely through interactions with quantitative sensory testing parameters and pain sensitivity; together, these parameters can explain a third of the pain variability irrespective of tissue damage. Fourth, these connectivity parameters do not change up to a year after the injury, and connectivity metrics collected at baseline, as well as the same parameters collected at six months, are able to predict subjects’ pain at six months after the initial injury. Together, these findings suggest an a-priori predisposition to pain grounded in brain WM properties, which results in higher pain ratings after acute injury. These findings shed light on brain mechanisms underlying pain sensitivity and inform the current literature of pain perception of a poorly understood patient population – mTBI patients.

Our study shows that an important part of the reported post-injury pain can be accounted for by brain imaging features alone. Although previous studies have successfully predicted experimental pain(15) and tonic pain(16) in healthy participants, as well as pain in patients at the chronic stage(16), this is, to the best of our knowledge, the first demonstration that early acute pain, isolated from plastic changes that occur over time after an injury, can be predicted based on brain structure. The strength of the evidence here lies in the clinical sample studied: these patients did not have any pain before injury and were scanned within hours of the accident, making them an ideal group to study injury-related, early acute pain in an ecological manner. The influence of the brain in pain perception is clear in the literature, but reports using brain structure and function to predict pain, particularly post-injury acute pain are scarce. Spisak and colleagues(17) have, for instance, recently showed that brain functional connectivity can predict pain sensitivity in healthy participants, which points to a predisposition to pain grounded on their underlying connectome(17). Here, we extend that idea by showing that injury-related pain can also be predicted from the structural connectome. Within mTBI/whiplash literature, there is an ongoing discussion about the origin of the reported pain: the majority of the time, it is difficult to find imaging correlates of structural damage(5–7), and the patient’s pain is affected by psychological aspects, such as psychological distress, anxiety, depression(8,18), and seem related to insurance pay-off and work-related disabilities(19). Mapping post-injury pain to brain measures of diffusivity solidifies, at least in part, the organic basis of the patients’ pain. The fact that the networks subtending this prediction are classically implicated in nociception (rather than more limbic brain regions implicated in negative affect), together with associations between these networks and psychophysical/psychological indices related to pain sensitivity, in particular, conditioned pain modulation, and pain sensitivity questionnaire scores, further adds that at this early time-point, pain sensitivity profiles—but not psychological aspects—dominate the association with pain ratings, which fits well with previous clinical findings(8).

Three WM connections reliably predicted pain in these patients: they appeared on all cross-validated folds, predicted pain within-sample and predicted pain in two additional independent groups of patients. The first, and strongest, predictive feature of post-injury pain was the number of WM connections between the primary motor and sensory regions of the brain. The implication of these two regions in pain perception is not novel, given that they are well-known components of nociceptive pathways(20). It is also well established in animal studies that the precentral and postcentral gyrus have reciprocal structural connections,(21,22) which are crucial for sensorimotor integration(23–28). In humans, imaging data supports the idea that there is strong information sharing between these two regions, both during task^30^ and during rest(30), which has been theorized to have a role in pain perception(31). The strength of connectivity in this area was also associated with conditioned pain modulation and pain sensitivity questionnaire scores, further suggesting that sensorimotor connectivity is related to pain sensitivity. This is also coherent with previous work showing that higher cortical density in the somatosensory cortex leads to less experienced pain while performing quantitative sensory tasks in healthy subjects(32) and that denser cortical thickness of the somatosensory cortex is associated with higher pain thresholds(33). In fact, it is known that the sensorimotor area is able to modulate pain, namely through direct projections to the thalamus(34). Our findings also show spatial specificity that coincide with the site of injury: connectivity between precentral and postcentral face and arm area (corresponding to the head/neck representation in the motor homunculus), as well as connectivity between the precentral face and postcentral trunk area (head/neck area in the sensory homunculus) are mostly driving these results.

Similarly, there is ample evidence supporting the association between the thalamus and pain as this region is the termination site of the spinothalamic tract and of nociceptive information relay to the cortex. The SPL is a multisensory, high-level integrative site for pain(35) and has also been associated with top-down modulation of pain(36). Connections between the thalamus and SPL have been described in tracing studies in rhesus monkeys, showing that the SPL projects to the ventrolateral (VL) and posterior lateral (VPL) thalamic nuclei(37). VL regions of the thalami are also connected to the parietal cortex in humans(13). This is precisely the thalamus region best predicting pain, as seen in the thalamic parcellation analysis. One recent study has shown that connections between the VL/VPL and the SPL (and the postcentral gyrus) are associated with the pathophysiology of pain(38), and higher gray matter density in the SPL has also been linked with pain sensitivity(32). Considering that SPL acts as a top-down inhibitor of sensory stimuli and given that thalamic-cortical projections from VPL to SPL are central to chronic pain, it is tempting to suggest that this connectivity may play a role in the inhibition (or exacerbation) of nociceptive signals arriving at the thalamus.

The third tract engaged in mTBI pain connects the planum polare to the superior lateral occipital cortex. The superior lateral occipital cortex (which in the Harvard-Oxford cortical atlas extends into the IPL, see Fig. 2C) is a main integral part of the default-mode network with ample evidence of manifesting in sensory and pain-related behavior(39). Its connection with the planum polare is surprising, as this region is traditionally involved in auditory processing(40). One critical aspect is that this region is located within an area rich in crossing-fibers (within the external capsule) and near important pain-related regions, such as the insula, the secondary somatosensory cortex, and the posterior operculum. Therefore, it is likely that the loss of clear directionality within the gray matter (a well-known limitation of DTI(41,42)) led to the fibers tractography algorithm to terminate prior to reaching the actual target. This explanation is speculative and requires future studies.

An important question to discuss is whether these brain features reflect injury-related parameters, short-term plastic changes, or rather an a-priori predisposition to pain hard-wired in the brain. Our results favor the latter. Considering an injury-related explanation to these results, one could argue these parameters are mapping tissue injury as a proxy for pain: it is known that whiplash-like centrifugal forces may cause axonal micro-lesions that are not detectable in conventional scanning protocols(43); and these brain lesions could, in turn, reflect how severe the mTBI/whiplash injury is(44), leading to higher reported pain. We find this possibility unlikely: first, we excluded patients with WAD scores above three and patients with obvious signs of brain damage. Second, we did not find associations between connectivity strength and whiplash severity, and the model was able to predict pain in the group of patients with no whiplash-like symptoms. Lastly, there are accounts that white-matter properties, namely fractional anisotropy and median diffusivity, are unaffected following mTBI(45). Another possibility is that this connectivity is reflecting short-term plastic changes caused by the new pain state. Our data does not support this hypothesis either: participants were scanned within 72h of the injury. It is unlikely that white-matter properties changed over a timespan of hours and ceased to change any further over longer timespans. There is evidence of short-term white-matter diffusivity changes in other contexts(46,47), but if the system is so malleable, one would expect to observe diffusivity changes over longer times too, especially as many participants gradually recover from their acute pain. We in fact observe no changes in connectivity strength in these WM networks from the time of the accident to six months and up to a year. Moreover, these same connections measured at six months post-injury also predict the reported pain at the same point in time, further adding strength to the within-subject reliability of this model. In conclusion, we consider it quite unlikely that our results reflect injury-related parameters or plastic changes that occur in the timespan of the data-collection; our data instead favors the idea that observed WM properties reflect an *a priori* brain predisposition for pain sensitivity.

While white-matter properties predicted pain quite successfully, fMRI functional connectivity did not. It is thought that functional connectivity is primarily supported by structural connectivity(48), and given the success of using resting-state to study pain sensitivity(17) and phasic pain(16), it is somewhat surprising that functional connectivity was not informative of acute mTBI pain. It is, however, important to point out key differences between DTI and rsfMRI techniques. While white-matter properties are relatively stable and should reflect primarily long-term changes in brain structure (i.e., more trait-like), rsfMRI should better reflect the emergence of a state within a given macro-structure (i.e., more state-like(49)). Given that patients were scanned within 72h of a motorized vehicle accident, an often emotionally charged event that will no doubt imprint a state of anxiety and distress on the patient, it may be difficult to identify a reliable and solid pattern that generalizes across subjects, particularly given the heterogeneity of the psychological factors and their influence on functional connectivity. Importantly, the fact that only white matter is predictive of pain post-injury in the patients should be construed as further evidence that they represent stable brain circuitry implicated in the transformation and amplification of nociceptive inputs, rather than simply capturing the perception (state) of pain per se.

This study has some limitations. We lack a group of healthy participants, which could be used to directly ascertain if the connectivity profiles are affected by the injury itself. Also, it is necessary to mention that the absolute agreement between predicted pain and observed pain is subpar (i.e. participants with 0 pain are predicted to have an average of 30 pain). Nonetheless, the goal of this study was to infer brain mechanisms from early acute pain, rather than minimizing the error in out-of-sample prediction. Finally, our sample is quite heterogenous in several respects, including acute treatment, type of car accident, and perhaps others. Naturally, there is an vast number confounds that could have been controlled for to improve the predictive ability of the model; here, we favored a less constrained approach as it provides strength to the predictive power of our analyses - it predicts pain regardless of uncontrolled confounds.

In conclusion, this study shows that measures of brain diffusivity in white-matter tracts predict an important part of acute pain shortly after an injury. These tracts are implicated on nociceptive and pain-related circuitry, which may reflect preexisting pro-nociceptive or anti-nociceptive influences that either amplify or diminish the cortical interpretation of the injury-related nociceptive barrage into the subjectivity of pain. These findings lead to new questions for future research: are these structural networks indeed generalizable across pain conditions? Can these connections predict acute, experimental pain in healthy subjects? The study of white-matter diffusivity in early acute pain thus opens new avenues for studying pain sensitivity and establishes dependence of pain perception, in part, on hard-wired central nervous system structures. It further provides support for the organic basis of pain perception in mTBI-whiplash patients, which is often labeled as purely psychogenic in nature.

## Materials and methods

### Participants

Participants included in this study were recruited at the Rambam Health Care Campus emergency department after suffering an mTBI injury from a motor vehicle accident at a maximum of 24 hours before their visit. They signed an informed consent in agreement with the Declaration of Helsinki. The study was approved by the institutional review board of Rambam Health Care Campus (ref. 0601-14). All patients had direct or indirect head and neck injury, a Glasgow Coma Scale score of 13 to 15 and no subsequent decline, and were over 18 years of age. We excluded patients with imaging-based traumatic brain findings and more than 30 minutes of unconsciousness. Patients were also excluded if they had other major bodily injuries, prior chronic head/neck pain, injuries in the head/neck area within one year prior to the current injury, and neurological diseases compromising the interpretation of brain function (e.g., neurogenerative diseases).

We recruited 249 patients, out of which we excluded 17 patients who did not fulfill mTBI clinical criteria, 54 patients who did not undergo MRI (claustrophobia, or not willing to participate), four patients who did not report baseline pain, and two patients with incidental MRI findings. Given the current discussion on the common etiology and symptomatology of patients with WAD and mTBI(50), we decided to homogenize our sample by studying only patients fulfilling both WAD and mTBI diagnosis (90% of patients). Hence, 15 patients showed no signs of WAD (WAD score = 0) and were excluded from the main analyses. They were, however, used post-hoc, as an additional validation group. The remaining 157 subjects (mean age = 37.3 ± 12 years, 86 male, average pain = 56.6 ± 26.2, range 0-100) were analyzed. Not all subjects completed all MRI protocols (detailed exclusions shown in Supplementary Fig. 1).

### Study design and procedures

After the initial emergency room visit, patients were informed about the study and were consented. Patients were then scheduled for a visit within 3 days of the injury (mean 1.7 days, range = 0–3), where they performed an MRI scan (structural, resting-state, and diffusion MRI protocols). Participants were asked to refrain from taking analgesic medication for 24 hours before the data collection. During the visit, patients were asked to rate their average pain over the previous 24 hours, separately, for their neck and head using a numeric rating scale (NRS, 0–100; 0: no pain, 100: worst imaginable pain); their current pain was determined to be the highest of the two. Patients also performed a series of clinical, psychological, and psychophysical tests, which are explained in detail elsewhere(8). Briefly, they filled a battery of psychological questionnaires measuring anxiety and depression (HADS(51)), pain catastrophizing (PCS(52)), pain sensitivity (PSQ(53)), stress (PSS(54)), and underwent quantitative sensory testing (conditioned pain modulation with heat and pressure stimuli, and temporal summation protocols with pressure and electrical stimuli). Some of these data are featured in other publications, examining psychophysical and psychological predictors of early acute mTBI pain(8,9).

### Scanner parameters

All images were collected on a 3T (MR 750, SIGNA 20, GE Medical Systems, Milwaukee, USA) scanner with a 16-channel head-coil. High-resolution T1-weighted images were collected with the following parameters: field of view (FOV) = 256 × 256 mm^2^, flip angle = 12°, slice thickness = 1 mm, in-plane pixel size 1 × 1 mm^2^, axial slices = 172. Diffusion weighted images were collected, with the following parameters: TR = 10000 ms, TE = 82, FOV = 256 × 256 mm^2^, slice thickness = 2 mm, in-plane pixel size 1 × 1 mm^2^, axial slices = 68, and number of directions = 60 with a b-value of 1000 s/ mm^2^. Five volumes with no diffusion weighting (b-value = 0 s/mm^2^) were acquired at the start of the protocol. Resting-state fMRI images were collected with an EPI sequence with the following parameters: TR = 2000 ms; TE = 30s; FOV = 220 × 220 mm^2^; flip angle = 75°; slice thickness = 3.4 mm; in-plane pixel size 3.44 × 3.44 mm^2^ and axial slices = 43. The first 8 s of acquisition in each run were excluded due to T1 equilibration effects. Participants were instructed to stay as still as possible, close their eyes, not fall asleep, and think of nothing in particular.

### Data preprocessing

All images were preprocessed using tools from the FMRIB FSL library (FSL 6.0.4(55)). Structural images were skull-stripped using BET(56) and segmented into white-matter, gray-matter, and cerebrospinal fluid (CSF) masks using FAST(57). As a proxy of total intracranial volume (TIV), the volume of grey-matter and white-matter masks were summed.

For resting-state fMRI images, data were skull-stripped, slice-time corrected, smoothed using a 6 mm FWHM kernel, and filtered using a band-pass filter (.02 to .001 Hz). To further remove signal of no interest, we used a strict denoising procedure: the six head motion parameters, their squared parameters, and the temporal derivatives of both were estimated. Signal from the WM and CSF was identified by averaging the time course within the respective masks; these masks were eroded once to ensure no overlap between tissue types. The 24 movement parameters, together with the WM and CSF time courses, were then regressed out from the data through multiple regression. Finally, high motion timepoints and their immediately adjacent volumes were removed (motion scrubbing) if they exceeded one of three criteria: timepoints with a framewise displacement larger than 0.7, timepoints exceeding 2.3 standard deviations of the mean signal, or a derivative of the root mean squared over voxels (i.e. DVARS) larger than 2.3. Functional images were normalized to standard MNI space using a two-step process. First, they were registered from functional to T1 structural space using boundary-based registration. Then, images were transformed from structural space to MNI by applying a 12 degrees-of-freedom linear transformation (FLIRT(58)), followed by a non-linear transformation (FNIRT). Subjects with less than 5 minutes of resting-state data after scrubbing were excluded (N = 3). Four additional subjects were excluded from analyses due to technical/image problems (see Supplementary Fig. 1). The final rsfMRI sample included 141 subjects.

Diffusion weighted images were visually inspected for obvious artefacts and corrected for eddy current distortions and head movement using EDDY(59). Then, a tensor was fit to the data, and principal directions were calculated using DTIFIT. To account for possible crossing-fibers, a Bayesian estimation of the principal directions was performed with BEDPOSTX(60), modelling up to three principal directions. Transformation matrices from subject space to MNI were obtained using FLIRT and FNIRT. For diffusion tensor imaging quality control, we used QUAD and SQUAD(61); subjects scoring as group outliers in absolute/relative displacement (N = 5) or signal to noise ratio/contrast to noise ratio (N = 1) were excluded. Eight other subjects were excluded due to technical issues with the data (see Supplementary Fig. 1). The final DTI sample includes 129 subjects.

### Extraction of brain connectivity features

To sample brain connectivity at a comprehensive yet interpretable scale, we used the Harvard-Oxford Cortical and Subcortical atlas. This atlas is composed of 115 brain regions and spans major brain networks. Due to field-of-view coverage and in line with our previous research, we excluded regions of interest (ROIs) within the cerebellum and the brainstem, resulting in a total of 105 cortical and subcortical ROIs (Supplementary Fig. 2A, B). For functional connectivity analyses, the average time course of each ROI was correlated to the time courses of all other ROIs, resulting in a 105 × 105 connectivity matrix. Since the matrix is symmetric, only the lower triangle of the connectivity matrix was kept, for a total of 5460 features per subject. Correlation (*r*) values were converted to z scores using the Fisher r-to-z conversion to approximate a normal distribution. For structural connectivity analyses, we first generated a white-matter/gray-matter interface mask by intersecting FSL’s white-matter and gray matter tissue priors at 25% probability (Supplementary Fig. 2C). Then, probabilistic tractography was ran using probtrackx2(60), seeding only from voxels within the ROIs that are included in the WM/GM interface mask. For each voxel within each ROI, 5000 streamlines were seeded, and the number of connections reaching each other ROI was counted, again resulting in a 105 × 105 WM connectivity matrix per subject. As distance between ROIs can influence connectivity results and introduce a head-size bias, WM connectivity count was corrected for distance by multiplying the number of streamlines by the Euclidian distance between the two ROIs (*--pd* option in probtrackx2). Finally, since the connectivity matrix was undirected, we averaged the upper and lower triangles of the matrix, to obtain a single ROI-to-ROI value indicative of probability of connectivity. As with rsfMRI connectivity, this resulted in 5460 structural connectivity features.

### Machine-learning approach

We built a machine-learning model to predict the patients’ reported pain, using whole-brain, structural and functional connectivity between 105 ROIs, yielding 5460 independent variables per method. To simplify this overdetermined fit – we studied 141 and 129 participants for rsfMRI and DTI, respectively – we used a well-established method, Connectome-based Predictive Modelling, CPM(62,63). This method simplifies the fit by reducing the number of features through a univariate feature selection, followed by a simplified form of coefficient regularization (see below). Univariate feature selection is performed by correlating all 5460 brain features against a dependent variable (Fig. 1A); these are then split into two sets of features, positively and negatively correlated to the dependent variable, which yield two separate models. Features correlating significantly below a given p-value are selected, which are summed to obtain one single parameter that characterizes the connectivity strength of a given set of tracts (Fig. 1B). Finally, the model is built by fitting a line between the dependent (pain) and independent variable (brain connectivity, Fig. 1C). Out of the 5460 features, this method reduces the fit to two coefficients (slope and intercept) per (positive and negative) model, which can then be tested for generalizability in the hold-out sample.

To enable model generalizability, we first split the dataset into two: a discovery dataset (70% of the data) on which the model was built and a hold-out dataset (30% of the data) on which the model was validated. This was performed using the Kennard-Stone algorithm(64), which divides a dataset into two or more partitions while matching them for given parameters, which we conducted based on age, sex, and TIV. Further, within the discovery dataset, we also performed a 10-fold cross-validation approach (Fig. 1D). The latter was needed for two goals: 1. To impartially determine the (otherwise arbitrary) univariate feature selection p-threshold; 2. To minimize model overfitting (more details in Supplementary Material). Thus, we divided the discovery dataset into 10 train and test splits and identified the p-threshold that best generalized within the discovery dataset (range of *p* = 0.001 to *p* = 0.1, p-value selected based on the best predictive performance, see Supplementary Fig. 3, 4). Finally, to generate a final model that can be tested in the hold-out sample and is consistent with other studies(62,65), we only kept features that appear on all of the CV folds; these were singled-out and refitted back to the whole discovery dataset (Fig. 1E). The resulting linear regression model was then used to generate predictions in the hold-out sample to assess generalizability (Fig. 1F). Performance metrics reported here are the correlation between predicted pain scores by the model and actual pain scores reported by the subject (*r*_predvsactual_). Functional and structural connectivity were modelled separately. As an additional quality control step, within the discovery and hold-out datasets, we exclude outliers – participants whose mean connectivity values are more than two standard deviations away from the group mean connectivity(66). This led to an exclusion of four participants from the rsfMRI dataset (all from the discovery dataset) and four participants from the DTI dataset (two from the discovery dataset, two from the hold-out dataset).

### Assessing generalizability of the machine-learning results

The above method can also be applied to less constrained models. To test the generalizability of the machine-learning pipeline under less stringent conditions, we tested the predictive ability of a simpler model by also performing univariate feature selection but instead feeding the selected features directly into a linear regression (i.e. not separating positive and negative correlations, and not summing all features into a single value). Again, in each CV-fold, significant features are selected and the threshold is determined by selecting the best performing model using the CV-samples. Models were validated like above. The same discovery and hold-out samples were used.

## Acknowledgements

The authors would like to thank all members of the Apkarian lab for their feedback on the manuscript. This work was supported by the Office of the Assistant Secretary of Defense for Health Affairs under award W81XWH-15-1-0603. Opinions, interpretations, conclusions, and recommendations are those of the authors and are not necessarily endorsed by the Department of Defense. This work was further supported by National Institutes of Health grant P50 DA044121 and grant R01AR074274-01A1.

